# C-terminal cysteines of HRas control Erk signaling and 15-deoxy-Δ^12,14^-prostaglandin J^2^ (15d-PGJ^2^) mediated inhibition of myoblast differentiation

**DOI:** 10.1101/2023.05.20.541575

**Authors:** Swarang Sachin Pundlik, Snehasudha Subhadarshini Sahoo, Alok Barik, Mahapatra Anshuman Jaysingh, Ashwin Venkateshvaran, Raviswamy G H Math, Arvind Ramanathan

**Affiliations:** Metabolic Regulation of Cell Fate (RCF), Institute for Stem Cell Science and Regenerative Medicine (InStem), Bangalore Life Science Cluster, GKVK - Post, Bellary Road, Bengaluru, Karnataka – 560065, India; Manipal Academy of Higher Education (MAHE), Tiger Circle Road, Madhav Nagar, Manipal, Karnataka – 576104, India; University of North Carolina at Chapel Hill, 216 Lenoir Dr, Chapel Hill, North Carolina – 27599 U. S. A; Department of Biological Sciences, Indian Institute of Science Education and Research Kolkata (IISER – K), Campus Road, Mohanpur, West Bengal – 741246, India; Division of Biology and Biomedical Sciences, Washington University in St Louis, 1 Brookings Dr, St. Louis, Missouri – 63130, U.S.A; National Centre for Biological Sciences (NCBS), Rajiv Gandhi Nagar, Kodigehalli, Bengaluru, Karnataka – 560065, India

## Abstract

HRas is an important node that controls cellular signaling, proliferation, and differentiation. Mutants of HRas (e.g., the constitutively active HRas V12) can be oncogenic, and can also inhibit myoblast differentiation. The C-terminal cysteines of HRas (Cys181 and Cys184) serve as substrates for intra-cellular reversible palmitoylation and de-palmitoylation reactions, which control its subcellular distribution. The relationship between the C-terminal cysteines of HRas, its intracellular distribution, and its cellular activity has remained unclear. Understanding this relationship has important implications for targeting HRas in pathogenic states where it is activated. In this study, we show that a mutation in the C-terminal of HRas, C181S, is sufficient to cause increased levels of HRas V12 in the Golgi, decreased HRas V12-driven Akt and Erk signaling and reverse the ability of HRas V12 to inhibit myoblast differentiation. This demonstrates the importance of C-terminal cysteines in controlling HRas V12. It has been previously shown that Cys184 can also be irreversibly modified by an electrophilic prostaglandin lipid 15d-PGJ_2_. This lipid is released by senescent cells as a part of senescence-associated secretory phenotype (SASP). In this study, we show that 15d-PGJ_2_ is secreted by senescent myoblasts formed by treatment with Doxorubicin. We also show that 15d-PGJ_2_ causes decreased levels of HRas within Golgi, activates Erk signaling (but not Akt signaling), and inhibits differentiation of C2C12 myoblasts in an HRas Cys184-dependent fashion. Chemotherapeutics such as Doxorubicin drive senescence and loss of skeletal muscle homeostasis in cancer patients. This study suggests that targeting the senescence-derived synthesis of 15-PGJ_2_ might be a target to promote muscle homeostasis after chemotherapy.

## Introduction

HRas belongs to the Ras superfamily of small 20 – 25 kDa GTPases(BOHR Orthopredic Hospital et al., 1964; Davis et al., 1983; Kirsten and Mayer, 1967; Vetter and Wittinghofer, 2001) and is a regulatory node for several key signaling pathways(Santra et al., 2019) regulating several cell biological processes. Activation of HRas is implicated in different types of cancers(Prior et al., 2012) and is known to transform immortalized non-malignant cells(Cheng et al., 2011). Constitutively active HRas has been shown to induce senescence(Bihani et al., 2007, 2004; Serrano et al., 1997). Constitutively activation of HRas signaling by overexpression of HRas V12 has been shown to inhibit the differentiation of myoblasts by inhibiting MyoD and Myogenin (MyoG) expression(Lassar et al., 1989; Liu et al., 2020; Olson,’ et al., 1987). Downstream signaling of HRas is important in maintaining muscle homeostasis, as constitutively active mutations in HRas (HRas V12) in patients suffering from RASopathies show skeletal and cardiac myopathies(Engler et al., 2021; Konieczny et al., 1989; Lee et al., 2010; Olson,’ et al., 1987; Scholz et al., 2009; Van Der Burgt et al., 2007). Post-translational modifications of the C-terminal hypervariable region regulate the intracellular distribution and activity of HRas. HRas undergoes reversible palmitoylation and de-palmitoylation at the C-terminal cysteines Cys181 and Cys184(Gutierrez et al., 1989; Lu and Hofmann, 1995), which regulate two spatial pools of HRas at the Golgi and the plasma membrane(Rocks et al., 2005). Downstream signaling of HRas involves two major pathways: the HRas – MAPK pathway and the HRas – PI3K pathway(Pylayeva-Gupta et al., 2011) which affect cell proliferation and metabolism respectively(Yu and Cui, 2016). The effect of intracellular distribution of constitutively active HRas on the MAPK and PI3K signaling pathways remains unclear. Previous studies suggest that differential availability of HRas effectors at different intracellular membranes(Santra et al., 2019). The effect of alterations in the intracellular distribution of HRas on cell physiology is also poorly understood. In this study, we demonstrate that the intracellular distribution of constitutively active HRas (HRas V12) differentially regulates HRas signaling in a C-terminal cysteine modification-dependent manner to regulate myoblast differentiation.

HRas has been previously shown to be covalently modified by the eicosanoid prostaglandin 15d-PGJ_2_, at Cys184(Luis Oliva et al., 2003). Covalent modification of HRas by 15d-PGJ_2_ has been shown to activate HRas, as seen by phosphorylation of Erk(Luis Oliva et al., 2003; Wiley et al., 2021). However, the effect of 15d-PGJ_2_ binding on HRas intracellular distribution has not been studied. We have previously shown that senescent lung fibroblast cells synthesize 15d-PGJ_2_ as a lipid component of (SASP)(Wiley et al., 2021). 15d-PGJ_2_ is a non-enzymatic dehydration product of prostaglandin PGD_2_(Shibata et al., 2002). PGD_2_ has been suggested to negatively regulate muscle differentiation independent of its cognate receptor(Hunter et al., 2001; Veliça et al., 2010). Therefore, inhibition of muscle differentiation by PGD_2_ could be via 15d-PGJ_2_. A cognate receptor for 15d-PGJ_2_ is yet to be identified, suggesting that a possible receptor-independent mechanism exists. In this study, we show that 15d-PGJ_2_ is secreted by senescent myoblasts which activates HRas signaling by the previously identified modification of C-terminal cysteine of HRas to inhibit differentiation of myoblasts.

## Results

### C181S mutant causes increased levels of EGFP-tagged HRas WT and HRas V12 at the Golgi compared to the plasma membrane

Palmitoylation and de-palmitoylation of two cysteine residues at the C-terminal of HRas (Cys181, Cys184) maintain two spatiotemporal pools of HRas localization and activity at the Golgi and the plasma membrane respectively(Rocks et al., 2005). We wanted to measure if the HRas GTPase activity somehow dictates HRas intracellular distribution. So, we transfected C2C12 mouse myoblasts with EGFP-tagged cysteine mutants of the wild type (HRas WT or HRas-C181S or HRas-C184S) or the constitutively active HRas (HRas V12 or HRas V12-C181S or HRas V12-C184S) and measured the distribution of HRas between the Golgi and the plasma membrane (Fig. 1A and Fig. S1A and B). We observed a significant increase in R_mean_, the ratio of mean HRas intensity at the Golgi to the mean HRas intensity at the plasma membrane, in the case of the C181S mutant (HRas-C181S and HRas V12-C181S) compared to the unmutated or the C184S mutant (HRas WT or HRas V12 and HRas-C184S or HRas V12-C184S) (Fig. 1B). This suggests that the acyl modifications of HRas at the C-terminal cysteines, predominantly at the Cys181, regulate the intracellular distribution of HRas independent of HRas GTPase activity.

**Figure 1:**
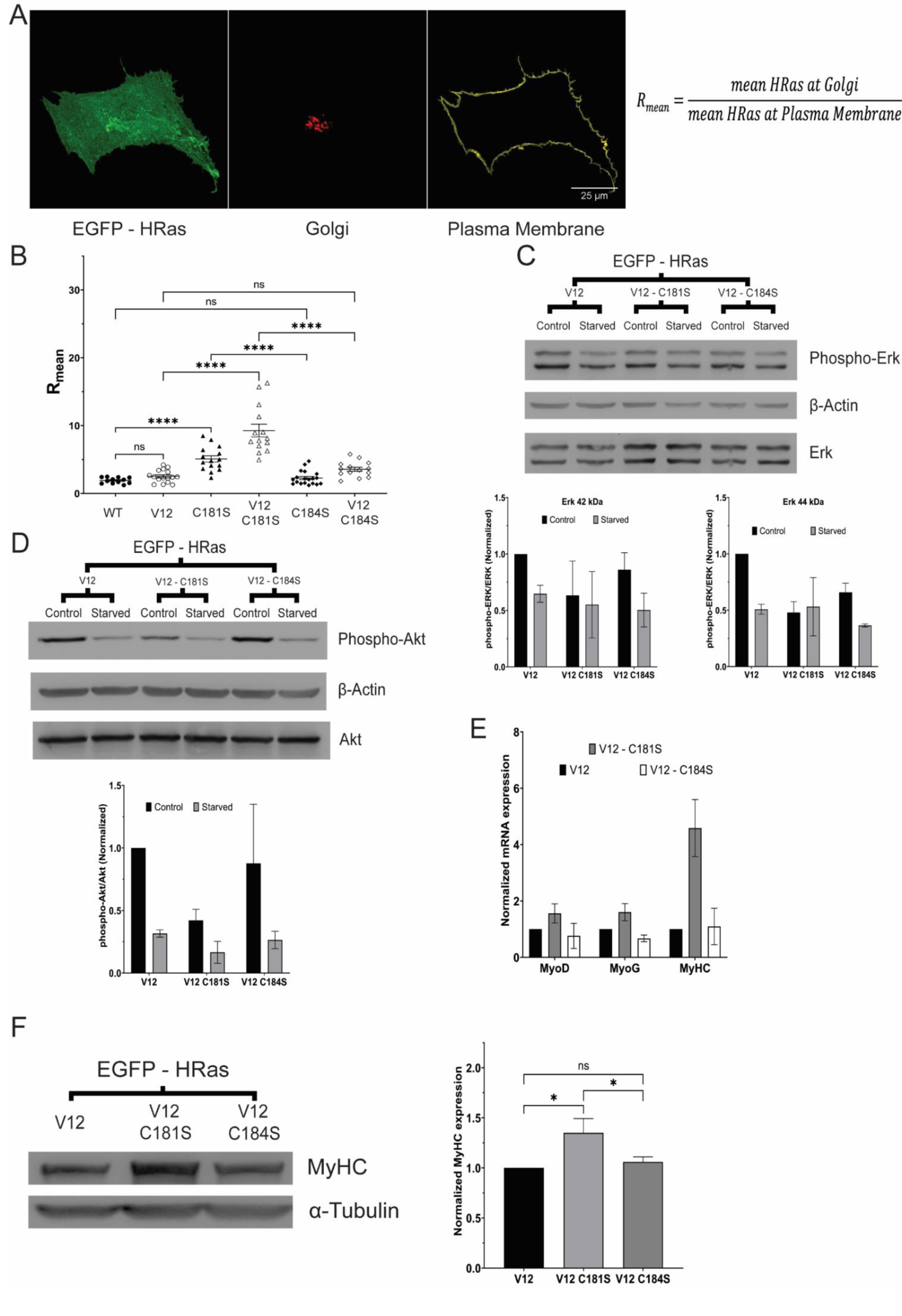
C-terminal cysteines mediated intracellular distribution and signaling of HRas V12 regulates differentiation of C2C12 myoblasts. A. A representative fluorescent confocal micrograph of C2C12 cell showing localization of HRas (EGFP-tagged) between the plasma membrane (stained with Alexa Fluor 633 conjugated Wheat Germ Agglutinin) and the Golgi complex (tagged with TagRFP conjugated Golgi resident GalT protein) B. Distribution of EGFP-tagged HRas WT/ HRas V12/ HRas-C181S/ HRas V12-C181S/ HRas-C184S/ HRas V12-C184S between the Golgi complex and the plasma membrane, as seen by the scatter plot of R_mean_, the ratio of the mean HRas intensity at the Golgi complex to that at the plasma membrane, in C2C12 cells. (Significance tested by two-tailed student’s t-test with the heteroscedastic distribution. ns = p>0.05, * = p<0.05, ** = p<0.01, *** = p<0.001, **** = p<0.0001) C. Phosphorylation of Erk1/2 (measured by immunoblotting) in C2C12 cells expressing EGFP-tagged HRas V12/ HRas V12-C181S/ HRas V12-C184S after 24 hours of starvation. The densitometric ratio of phosphorylated Erk1/2 (Thr202/Tyr204) to total Erk1/2 was normalized to non-starved C2C12 cells. (Significance tested by two-tailed student’s t-test with the heteroscedastic distribution. ns = p>0.05, * = p<0.05, ** = p<0.01, *** = p<0.001, **** = p<0.0001) D. Phosphorylation of Akt (measured by immunoblotting) in C2C12 cells expressing EGFP-tagged HRas V12/ HRas V12-C181S/ HRas V12-C184S after 24 hours of starvation. The densitometric ratio of phosphorylated Akt (Ser473) to total Akt was normalized to non-starved C2C12 cells. (Significance tested by two-tailed student’s t-test with the heteroscedastic distribution. ns = p>0.05, * = p<0.05, ** = p<0.01, *** = p<0.001, **** = p<0.0001) E. mRNA levels of MyoD, MyoG, and MyHC relative to 18s rRNA (measured by quantitative PCR) in C2C12 cells expressing EGFP-tagged HRas V12/ HRas V12-C181S/ HRas V12-C184S in C2C12 differentiation medium for 5 days. (Significance tested by two-tailed student’s t-test with the heteroscedastic distribution. ns = p>0.05, * = p<0.05, ** = p<0.01, *** = p<0.001, **** = p<0.0001) F. Protein levels of MyHC (measured by immunoblotting) in C2C12 cells expressing EGFP-tagged HRas V12/HRas V12-C181S/HRas V12-C184S in C2C12 differentiation medium for 5 days. The densitometric ratio of levels of MyHC to α-Tubulin was normalized to C2C12 cells expressing HRas V12. (Significance tested by two-tailed student’s t-test with the heteroscedastic distribution. ns = p>0.05, * = p<0.05, ** = p<0.01, *** = p<0.001, **** = p<0.0001)

### C181S mutant decreases HRas V12 mediated phosphorylation of Erk and Akt

We wanted to measure the effect of the intracellular distribution of HRas between the Golgi and the plasma membrane on downstream signaling. HRas regulates two major downstream signaling pathways, the MAP kinase (MAPK) pathway and the PI3 kinase (PI3K) pathway(Pylayeva-Gupta et al., 2011). So, we measured the phosphorylation of Erk (42 kDa and 44 kDa) proteins and Akt protein, key mediators in the MAPK signaling and PI3K signaling respectively, in C2C12 mouse myoblasts expressing EGFP-tagged HRas V12 or HRas V12-C181S or HRas V12-C184S. We observed a decrease in the phosphorylation of Erk (42 kDa) and Akt in C2C12 cells expressing the HRas V12-C181S but not HRas V12-C184S compared to C2C12 cells expressing HRas V12 (Fig. 1C and D). We also observed a decrease in the phosphorylation of Erk (44 kDa) protein in C2C12 cells expressing both HRas V12-C181S and HRas V12-C184S. These observations suggest diminished HRas downstream signaling by HRas V12-C181S, which is predominantly localized at the Golgi. Then, we measured the effect of HRas intracellular distribution on changes in HRas downstream signaling upon serum starvation. We observed a decrease in phosphorylation of both Erk (42 kDa and 44 kDa) in C2C12 cells expressing HRas V12 and HRas V12-C184S upon serum starvation but not in C2C12 cells expressing HRas V12-C181S (Fig. 1C). However, we observed a decrease in Akt phosphorylation in C2C12 cells expressing EGFP-HRas V12, EGFP-HRas V12-C181S, and EGFP-HRas V12-C184S upon serum starvation (Fig. 1D). These observations suggest that HRas-PI3K signaling responds to serum starvation, but not HRas-MAPK signaling at the Golgi.

### C181S mutant rescues the HRas V12 mediated inhibition of myoblast differentiation

Constitutively active HRas (HRas V12) is a known inhibitor of differentiation in myoblasts(Konieczny et al., 1989; Lee et al., 2010; Olson,’ et al., 1987; Scholz et al., 2009; Van Der Burgt et al., 2007). Therefore, we wanted to examine the effect of altered intracellular distribution and downstream signaling of HRas in cysteine mutants of HRas V12 on differentiation of C2C12 myoblasts. We measured the mRNA levels of differentiation markers MyoD, MyoG, and MyHC in C2C12 cells expressing EGFP-tagged HRas V12 or HRas V12-C181S or HRas V12-C184S. We observed an increase in mRNA levels of MyoD, MyoG, and MyHC in differentiating C2C12 cells expressing HRas V12-C181S but not HRas V12-C184S compared to C2C12 expressing HRas V12 (Fig. 1E). We also observed an increase in the protein levels of MyHC in differentiating C2C12 myoblasts expressing HRas V12-C181S but not HRas V12-C184S compared to C2C12 expressing HRas V12 (Fig. 1F). This suggests altered HRas downstream signaling rescues inhibition of differentiation in C2C12 myoblasts (Fig. S1C)

### Electrophilic lipid 15d-PGJ_2_ causes decreased levels of EGFP-tagged HRas WT at the Golgi compared to the plasma membrane

15d-PGJ_2_ is a non-enzymatic dehydration product of prostaglandin PGD_2_(Shibata et al., 2002) and is hypothesized to inhibit muscle differentiation(Hunter et al., 2001). Also, 15d-PGJ_2_ is shown to covalently modify HRas(Luis Oliva et al., 2003). So, we wanted to measure the effect of 15d-PGJ_2_ binding on the intracellular distribution of HRas. We measured the distribution of HRas between the Golgi and the plasma membrane in C2C12 cells expressing EGFP-tagged HRas WT after 15d-PGJ_2_ (10 µM) treatment. We observed a decrease in R_mean_ in C2C12 cells expressing EGFP-tagged HRas WT after 24 hrs of 15d-PGJ_2_ treatment (Fig. 2A), suggesting the redistribution of HRas from the Golgi to the plasma membrane after 15d-PGJ_2_ treatment.

**Figure 2:**
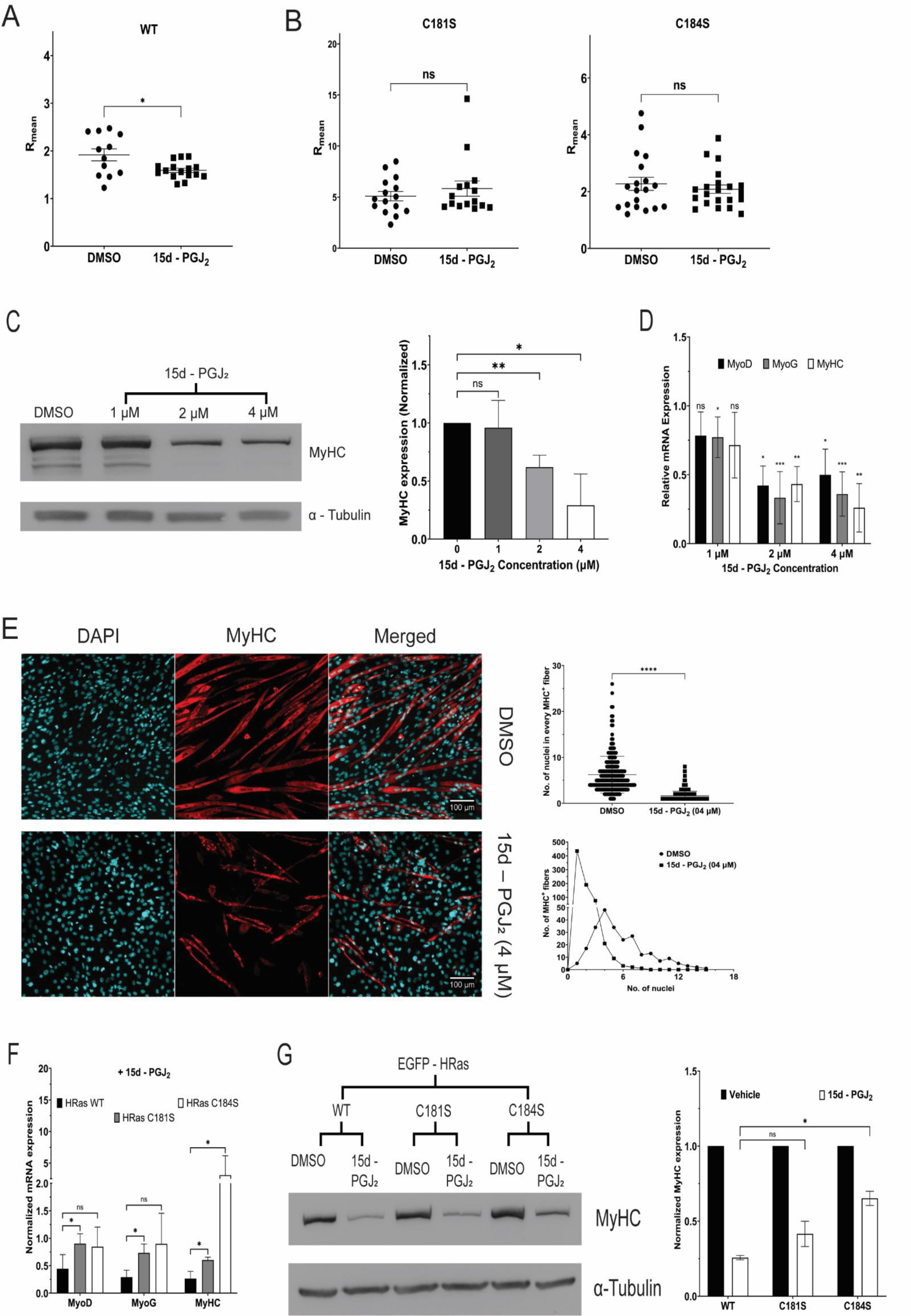
15d-PGJ_2_ decreases levels of HRas in the Golgi compared to the plasma membrane in C2C12 cells and inhibits differentiation in an HRas C-terminal cysteine mutation-dependent manner. A. Distribution of EGFP-tagged HRas WT between the Golgi complex and the plasma membrane, as seen by the scatter plot of R_mean_, the ratio of the mean HRas intensity at the Golgi complex to that at the plasma membrane, in C2C12 cells treated with 15d-PGJ_2_ (10 µM) or DMSO for 24 hrs. (Significance tested by two-tailed student’s t-test with the heteroscedastic distribution. ns = p>0.05, * = p<0.05, ** = p<0.01, *** = p<0.001, **** = p<0.0001) B. Distribution of EGFP-tagged HRas-C181S and HRas-C184S between the Golgi complex and the plasma membrane, as seen by the scatter plot of the ratio of the mean HRas intensity at the Golgi complex to that at the plasma membrane, in C2C12 cells treated with 15d-PGJ_2_ (10 µM) or DMSO for 24 hrs. (Significance tested by two-tailed student’s t-test with the heteroscedastic distribution. ns = p>0.05, * = p<0.05, ** = p<0.01, *** = p<0.001, **** = p<0.0001) C. Protein levels of MyHC (measured by immunoblotting) in differentiating primary human myoblasts treated with 15d-PGJ_2_ (1, 2, 4 µM) for 5 days. The densitometric ratio of MyHC levels relative to α-Tubulin was normalized to Human myoblasts treated with DMSO. (Significance tested by two-tailed student’s t-test with the heteroscedastic distribution. ns = p>0.05, * = p<0.05, ** = p<0.01, *** = p<0.001, **** = p<0.0001) D. mRNA levels of MyoD, MyoG, and MyHC relative to 18s rRNA (measured by quantitative PCR) in C2C12 cells treated with 15d-PGJ_2_ (1, 2, 4 µM) for 5 days in C2C12 differentiation medium compared those treated with DMSO. (Significance tested by two-tailed student’s t-test with the heteroscedastic distribution. ns = p>0.05, * = p<0.05, ** = p<0.01, *** = p<0.001, **** = p<0.0001) E. Fusion of myoblasts into myotubes, measured by immunofluorescence followed by a scatter plot and histogram of the number of nuclei per MHC^+^ fiber, in differentiating C2C12 cells treated with 15d-PGJ_2_ (4 µM) or DMSO for 5 days in C2C12 differentiation medium. (Significance tested by two-tailed student’s t-test with the heteroscedastic distribution. ns = p>0.05, * = p<0.05, ** = p<0.01, *** = p<0.001, **** = p<0.0001) F. mRNA levels of MyoD, MyoG, and MyHC relative to 18s rRNA (measured by quantitative PCR) in C2C12 cells expressing EGFP-tagged HRas WT/ HRas-C181S/ HRas-C184S in C2C12 cells treated with 15d-PGJ_2_ (4 µM) or DMSO in C2C12 differentiation medium for 5 days. (Significance tested by two-tailed student’s t-test with the heteroscedastic distribution. ns = p>0.05, * = p<0.05, ** = p<0.01, *** = p<0.001, **** = p<0.0001) G. Protein levels of MyHC (measured by immunoblotting) in C2C12 cells expressing EGFP-tagged HRas WT/ HRas-C181S/ HRas-C184S in C2C12 cells treated with 15d-PGJ_2_ (4 µM) or DMSO in C2C12 differentiation medium for 5 days. The densitometric ratio of MyHC levels relative to α-Tubulin was normalized to C2C12 myoblasts treated with DMSO. (Significance tested by two-tailed student’s t-test with the heteroscedastic distribution. ns = p>0.05, * = p<0.05, ** = p<0.01, *** = p<0.001, **** = p<0.0001)

### Electrophilic lipid 15d-PGJ_2_ does not change the levels of EGFP-tagged HRas C181S and HRas C184S mutants at the Golgi compared to the plasma membrane

15d-PGJ_2_ is shown to covalently modify HRas at Cys184(Luis Oliva et al., 2003). So, we wanted to examine the role of the Cys181 and Cys184 in 15d-PGJ_2_ mediated redistribution of HRas. We measured the distribution of HRas between the Golgi and the plasma membrane in C2C12 cells expressing EGFP-tagged HRas-C181S or HRas-C184S after 15d-PGJ_2_ (10 µM) treatment. We did not observe a decrease in R_mean_ in C2C12 cells expressing HRas-C181S and HRas-C184S after 15d-PGJ_2_ treatment (Fig. 2B). This suggests 15d-PGJ_2_ mediated redistribution of HRas requires C-terminal cysteines.

### 15d-PGJ2 inhibits the differentiation of C2C12 and primary human myoblasts

Inhibition of myoblast differentiation by prostaglandin PGD_2_ has been hypothesized to cause by 15d-PGJ_2_(Veliça et al., 2010). So, we measured the differentiation of myoblasts after 15d-PGJ_2_ treatment. In differentiating medium, we observed cell death upon treating C2C12 myoblasts with 15d-PGJ2 (5 µM and 10 µM) (Fig. S2A). So, we treated the myoblasts with 15d-PGJ_2_ (1 µM/ 2 µM/ 4 µM). We observed a significant decrease in protein levels of MyHC in primary human myoblasts after 15d-PGJ_2_ treatment in a dose-dependent manner (Fig. 2C). We also observed a decrease in mRNA levels of MyoD, MyoG, and MyHC after 15d-PGJ_2_ treatment in a dose-dependent manner (Fig. 2D). We also observed a decrease in the fusion of myoblasts to myotubes in C2C12 treated with 15d-PGJ_2_ (4 µM) compared to DMSO (Fig. 2E). These observations suggest 15d-PGJ_2_ inhibits differentiation of myoblasts.

### 15d-PGJ2 mediated inhibition of C2C12 myoblast differentiation is rescued by HRas C181S and C184S mutants

15d-PGJ_2_ has been shown to covalently bind to HRas(Luis Oliva et al., 2003), we wanted to examine the role of HRas-15d-PGJ_2_ interaction on inhibition of myoblast differentiation by 15d-PGJ_2_. We treated C2C12 cells expressing EGFP-tagged HRas WT or HRas-C181S or HRas-C184S with 15d-PGJ_2_ (4 µM). We observed a decrease in mRNA levels of MyoD, MyoG, and MyHC in C2C12 cells expressing HRas WT after 15d-PGJ_2_ treatment, which was absent from C2C12 cells expressing HRas-C181S and HRas-C184S (Fig. 2F). We also observed a decrease in proteins levels of MyHC in C2C12 cells expressing HRas WT after 15d-PGJ_2_ (4 µM) treatment, which was partially rescued in C2C12 cells expressing HRas-C181S and HRas-C184S (Fig. 2G). This suggests the inhibition of myoblast differentiation by 15d-PGJ_2_ involves HRas C-terminal cysteines, predominantly Cys184.

### 15d-PGJ2 increases phosphorylation of Erk (Thr202/Tyr204) but not Akt (S473) in C2C12 myoblasts

We wanted to measure the effect of altered intracellular distribution of HRas after 15d-PGJ_2_ treatment on the downstream signaling. So, we measured the phosphorylation of Erk (42 kDa and 44 kDa) and Akt proteins in C2C12 cells after 15d-PGJ_2_ treatment. We treated C2C12 cells with 15d-PGJ_2_ (5 µM and 10 µM) or DMSO for 1 hr after starving the cells for 24 hrs, and observed a dose-dependent increase in the phosphorylation of Erk (42 kDa) but not of Erk (44 kDa) (Fig. 3A). we did not observe an increase in the phosphorylation of Akt in C2C12 cells after 15d-PGJ_2_ treatment (Fig. 3B). This suggests activation of MAPK signaling pathway but not PI3K signaling pathway by 15d-PGJ_2_.

**Figure 3:**
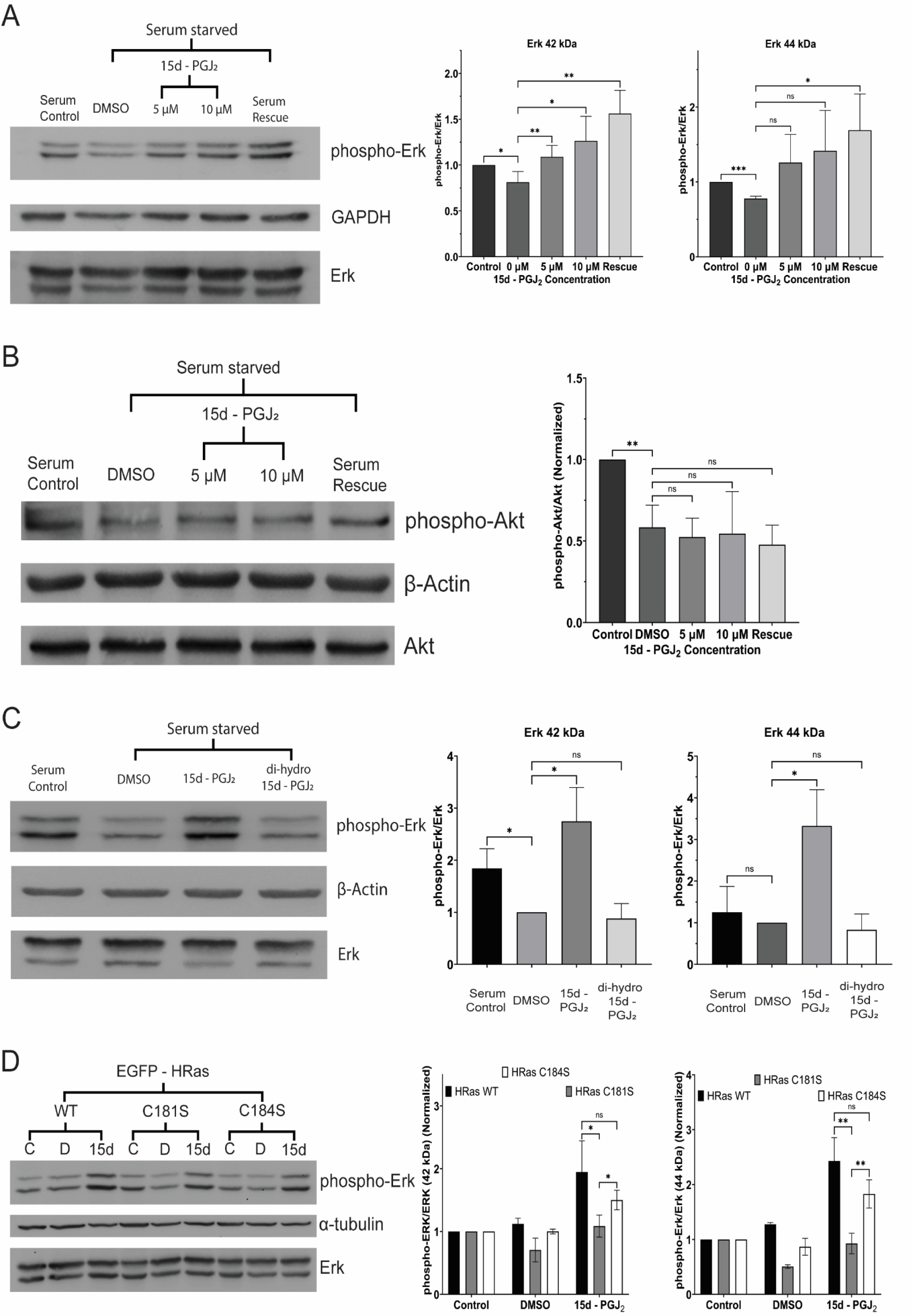
15d-PGJ_2_ activates MAPK but not PI3K signaling via the electrophilic cyclopentenone ring in an HRas C-terminal cysteine mutation-dependent manner. A. Phosphorylation of Erk (42 kDa and 44 kDa) (measured by immunoblotting) in C2C12 cells treated with 15d-PGJ_2_ (5, 10 µM) or DMSO for 1 hr after starving the cells in 0.2% serum medium for 24 hrs. The densitometric ratio of phosphorylated Erk (Thr202/ Tyr204) to total Erk was normalized to non-starved C2C12 cells. (Significance tested by two-tailed student’s t-test with the heteroscedastic distribution. ns = p>0.05, * = p<0.05, ** = p<0.01, *** = p<0.001, **** = p<0.0001) B. Phosphorylation of Akt (measured by immunoblotting) in C2C12 cells treated with 15d-PGJ_2_ (5, 10 µM) or DMSO for 1 hr after starving the cells in 0.2% serum medium for 24 hrs. The densitometric ratio of phosphorylated Akt (Ser473) to total Akt was normalized to non-starved C2C12 cells. (Significance tested by two-tailed student’s t-test with the heteroscedastic distribution. ns = p>0.05, * = p<0.05, ** = p<0.01, *** = p<0.001, **** = p<0.0001) C. Phosphorylation of Erk (42 kDa and 44 kDa) (measured by immunoblotting) in C2C12 cells treated with 15d-PGJ_2_ (10 µM)/ 9,10-dihydro-15d-PGJ_2_ (10 µM) or DMSO for 1 hr after starving the cells in 0.2% serum medium for 24 hrs. The densitometric ratio of phosphorylated Erk (Thr202/ Tyr204) to total Erk was normalized to non-starved C2C12 cells. (Significance tested by two-tailed student’s t-test with the heteroscedastic distribution. ns = p>0.05, * = p<0.05, ** = p<0.01, *** = p<0.001, **** = p<0.0001) D. Phosphorylation of Erk (42 kDa and 44 kDa) (measured by immunoblotting) C2C12 cells expressing EGFP-tagged HRas WT/ HRas-C181S/ HRas-C184S and treated with 15d-PGJ_2_ (10 µM) or DMSO for 1 hr after starving the cells in 0.2% serum medium for 24 hrs. The densitometric ratio of phosphorylated Erk (Thr202/ Tyr204) to total Erk was normalized to non-starved C2C12 cells. (Significance tested by one-tailed student’s t-test with the heteroscedastic distribution. ns = p>0.05, * = p<0.05, ** = p<0.01, *** = p<0.001, **** = p<0.0001)

### 9,10-dihydro-15d-PGJ2, which lacks a reactive electrophilic center, does not increase the phosphorylation of Erk (Thr202/Tyr204)

While a cognate receptor for 15d-PGJ_2_ is not yet identified, 15d-PGJ_2_ is previously shown to act via the electrophilic center in the cyclopentenone ring(Luis Oliva et al., 2003). So, we measured the role of the electrophilic center in the cyclopentenone ring in activation of MAPK signaling pathway. We measured the phosphorylation of Erk (42kDa and 44 kDa) after treating the cells with 9,10-dihydro-15d-PGJ_2_ (10 µM), a 15d-PGJ_2_ analog devoid of the electrophilic center, for 1 hr after starving the cells for 24 hrs. We observed the abrogation of 15d-PGJ_2_ mediated phosphorylation of Erk (42 kDa and 44 kDa) in C2C12 cells treated with 9,10-dihydro-15d-PGJ_2_ (Fig. 3C). This suggests that activation of MAPK signaling pathway by 15d-PGJ_2_ involves the electrophilic center in the 15d-PGJ_2_ cyclopentenone ring.

### 15d-PGJ2 mediated phosphorylation of Erk (Thr202/ Tyr204) is inhibited in C2C12 myoblasts expressing EGFP-tagged HRas C181S

15d-PGJ_2_ has been previously shown to covalently modify HRas at Cys184(Luis Oliva et al., 2003). Therefore, we measured the phosphorylation of Erk (42 kDa and 44 kDa) in C2C12 cells expressing the cysteine mutants of HRas after 15d-PGJ_2_ treatment. We treated C2C12 cells expressing EGFP-tagged HRas WT or HRas-C181S or HRas-C184S with 15d-PGJ_2_ (10 µM) for 1hr (after starvation for 24 hrs). We observed an increase in phosphorylation of Erk (both 42kDa and 44 kDa) (Thr202/ Tyr204) in C2C12 cells expressing HRas WT or HRas-C184S, but not in C2C12 cells expressing HRas-C181S (Fig. 3D). We also observed that the increase in the phosphorylation of Erk is decreased in C2C12 cells expressing HRas-C184S compared to C2C12 cells expressing HRas WT (Fig. 3D). This suggests a differential role of the two HRas C-terminal cysteines in 15d-PGJ_2_ mediated phosphorylation of Erk.

### DNA damage-induced senescence induces the synthesis of 15d-PGJ_2_ in mice muscles and C2C12 myoblasts

Synthesis and release of prostaglandins as SASP factors by senescent cells has previously been reported (Wiley and Campisi, 2021). Therefore, we measured the levels of 15d-PGJ_2_ synthesized and secreted in C2C12 myoblasts after induction of senescence by Doxorubicin (Doxo)-induced DNA damage. We treated C2C12 myoblasts with Doxo (150 nM) and observed flattened morphology of C2C12 cells treated with Doxo but not that with DMSO (Fig. S4A). We also observed an increase in the size of Nuclei of C2C12 cells treated with Doxo but not with DMSO (Fig. S4B) and increased senescence-associated β galactosidase (SA-β gal) activity in cells treated with Doxo but not with DMSO (Fig. S4C). This shows the induction of senescence in C2C12 cells after treatment with Doxo. We detected a significant increase in the levels of 15d-PGJ_2_ in the conditioned medium of senescent C2C12 myoblasts as compared to non-senescent myoblasts (Fig. 4A). We detected a significant increase in the intracellular levels of 15d-PGJ_2_ in senescent C2C12 myoblasts (Fig. 4B). These observations show an increase in the synthesis and release of 15d-PGJ_2_ by senescent C2C12 cells.

**Figure 4:**
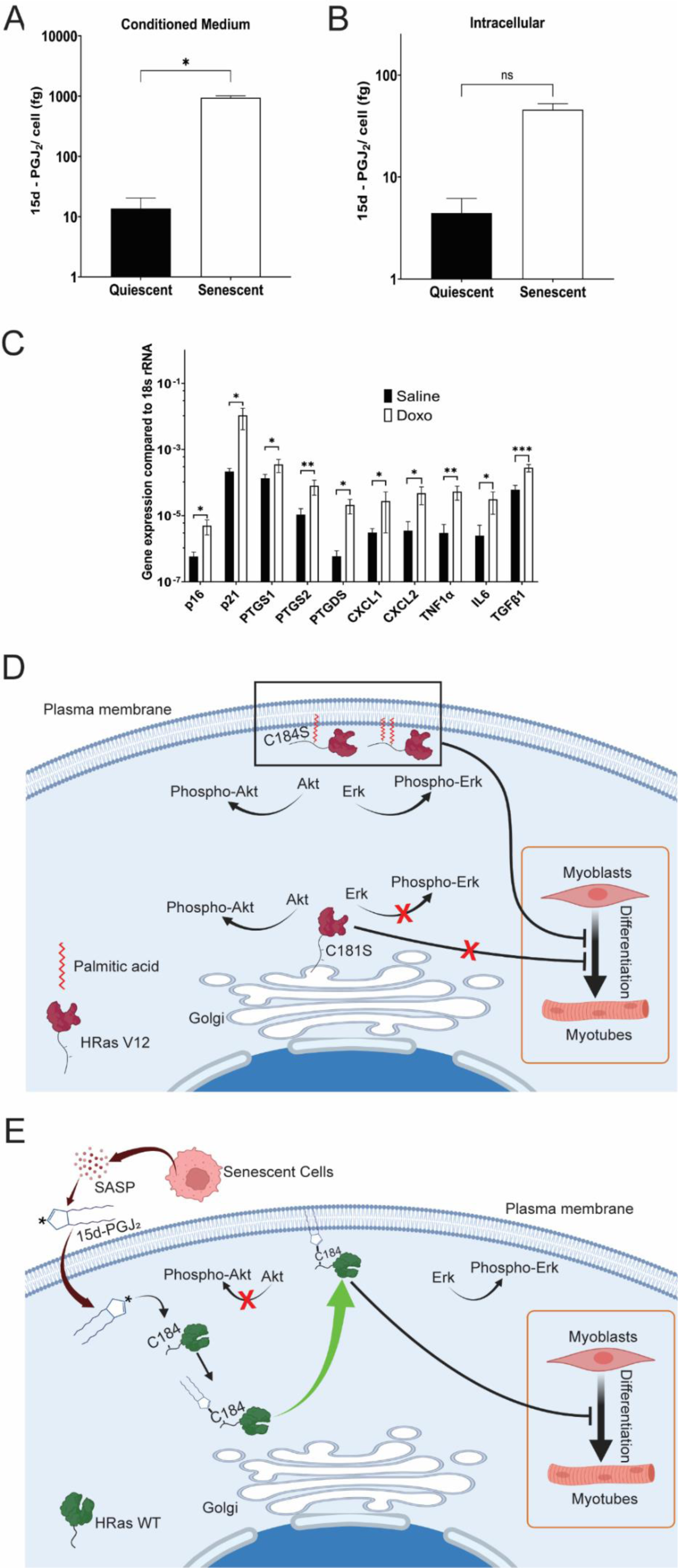
15d-PGJ_2_ synthesis and secretion are upregulated after Doxorubicin (Doxo)-mediated DNA damage-induced senescence. A. Concentration of 15d-PGJ_2_ in the conditioned medium of C2C12 myoblasts treated with Doxorubicin (150 nM) (Senescent) or DMSO (Quiescent). (Significance tested by two-tailed student’s t-test with the heteroscedastic distribution. ns = p>0.05, * = p<0.05, ** = p<0.01, *** = p<0.001, **** = p<0.0001) B. Intracellular concentration of 15d-PGJ_2_ in C2C12 myoblasts treated with Doxorubicin (150 nM) (Senescent) or DMSO (Quiescent). (Significance tested by two-tailed student’s t-test with the heteroscedastic distribution. ns = p>0.05, * = p<0.05, ** = p<0.01, *** = p<0.001, **** = p<0.0001) C. mRNA levels of markers of senescence (p16 and p21), SASP factors (CXCL1, CXCL2, TNF1α, IL6, TGFβ1), and enzymes involved in prostaglandin PGD_2_/15d-PGJ_2_ biosynthesis (PTGS1, PTGS2, PTGDS) (measured by quantitative PCR) in skeletal muscle of mice after 11 days after treating with Doxorubicin (5mg/kg) or Saline (treated every 3 days). (Significance tested by two-tailed student’s t-test with the heteroscedastic distribution. ns = p>0.05, * = p<0.05, ** = p<0.01, *** = p<0.001, **** = p<0.0001) D. A graphical summary of cellular mechanism showing regulation of intracellular HRas signaling and differentiation of myoblasts by C-terminal cysteine-dependent intracellular distribution of HRas V12. E. A graphical summary of mechanism showing regulation of HRas intracellular signaling by C-terminal cysteines during 15d-PGJ_2_ mediated inhibition of myoblast.

We next measured the induction of senescence and the levels of enzymes involved in the synthesis of PGD_2_/15d-PGJ_2_ in the skeletal muscles after treatment with Doxo. We found an increase in the mRNA levels of known senescence markers, p16 and p21 and known SASP factors: IL6, TNF1α, TGFβ1, CXCL1, and CXCL2 in the gastrocnemius muscle of mice after treatment with Doxo (Fig. 4C). We also observed increased nuclear levels of γH2A.X and p21 in the sections of the gastrocnemius muscle of mice treated with Doxo as compared to those treated with saline (Fig. S4E and F), suggesting the induction of senescence in mice skeletal muscle. We measured the mRNA levels of enzymes involved in the synthesis of PGD_2_ (PTGS1, PTGS2, and PTGDS), in the gastrocnemius muscle. We observed an increase in the mRNA levels of PTGS1, PTGS2, and PTGDS enzymes in skeletal muscle of mice treated with Doxo (Fig. 4C). These observations suggest that pathways involved in the synthesis of PGD_2_ and 15d-PGJ_2_ are activated in mice after induction of senescence.

## Discussion

HRas is one of the three major isoforms in the Ras family of small molecular weight GTPases in humans(BOHR Orthopredic Hospital et al., 1964; Davis et al., 1983; Kirsten and Mayer, 1967; Vetter and Wittinghofer, 2001). Point mutations in the *ras* gene constitutively activate the Ras protein(Reddy et al., 1982), which is found in 30% of solid tumors(Bos, 1989; Prior et al., 2012). Conformational changes in the GTPase domain of HRas regulate the activity of HRas, and mutations in the G domain (G12V) can activate HRas constitutively (HRas V12)(Reddy et al., 1982). Post-translational palmitoylation of Cysteines in the C-terminal tail of HRas (Cys181, Cys184) regulate the subcellular localization of HRas between intracellular membranes(Gutierrez et al., 1989; Lu and Hofmann, 1995; Rocks et al., 2005). Palmitoylation and de-palmitoylation of C-terminal cysteines maintain two spatially distinct pools of HRas at the Golgi and the plasma membrane respectively(Rocks et al., 2005). However, the effect of the post-translational acylations on the HRas signaling and its cellular activity is not well understood. Constitutively active HRas has been shown to induce senescence in several cells like MCF7, MDA-MB 231, IMR90 MEFs, etc.(Bihani et al., 2007, 2004; Serrano et al., 1997). Constitutively activation of HRas signaling by overexpression of HRas V12 has been shown to inhibit the differentiation of myoblasts, primarily by inhibition of MyoD and Myogenin (MyoG) expression(Lassar et al., 1989; Liu et al., 2020; Olson,’ et al., 1987). In this study, we show that mutations in the C-terminal cysteines affect the intracellular signaling of HRas V12 (Fig. 4D). The C181S mutant of HRas V12 (HRas V12-C181S) is predominantly localized to the Golgi as compared to the C184 mutant (HRas V12-C184S) or HRas V12 (HRas V12) (Fig. 1B). C2C12 cells expressing HRas V12-C181S show decreased Erk and Akt phosphorylation compared to C2C12 expressing HRas V12-C184S or HRas V12 (Fig. 1C and D). Erk and Akt are key mediators of the MAPK and the PI3K signaling pathways respectively. HRas regulates both the MAPK and the PI3K pathway(Pylayeva-Gupta et al., 2011). Therefore, our findings suggest that the intracellular distribution of constitutively active HRas affects intracellular signaling. We also show that mutations in the C-terminal cysteines regulate HRas V12-mediated inhibition of differentiation of myoblasts into myoblasts (Fig. 4D). C2C12 myoblasts expressing HRas V12-C181S differentiate into myotubes while those expressing HRas V12 or HRas V12-C184S fail to do so (Fig. 1E). This suggests that Cys181 controls intracellular distribution and signaling of HRas V12 and inhibition of differentiation of myoblast into myotubes. This relationship between the activity of HRas V12 and its sub-cellular localization provides new insights into the regulation of HRas signaling. Therefore, targeting the subcellular localization of HRas via Cys181 could be a possible strategy for controlling the activity of HRas V12.

Prostaglandins are naturally occurring metabolites of Arachidonic Acid (AA)(Lands, 1979). Prostaglandins have been shown to regulate several aspects of muscle regeneration. Prostaglandin degrading enzyme 15PGDH has been shown to be upregulated in aged murine muscles, and the inhibition of this enzyme has been shown to reverse sarcopenia in a PGE_2_-dependent manner(Palla et al., 2021). Mechanical stretch-dependent differentiation of myoblasts is shown to be driven by the activity of PTGS2, the enzyme involved in the initial steps of prostaglandin biosynthesis(Otis et al., 2005). Exercise-induced increase in differentiation of myoblasts has been shown to be dependent on PGE_2_ and PGF_2α_ levels(Trappe et al., 2001). On the other hand, prostaglandin PGD_2_ has been shown to inhibit myoblast differentiation in a receptor-independent manner(Veliça et al., 2010). However, the mechanism behind the inhibition of muscle differentiation by PGD_2_ remains unknown. In this study, we show that 15d-PGJ_2_, a non-enzymatic metabolite of PGD_2_(Shibata et al., 2002), inhibits the differentiation of myoblasts by regulation of HRas signaling (Fig. 4E). We show that 15d-PGJ_2_ inhibits differentiation of C2C12 myoblasts expressing the wild type HRas. However, inhibition of differentiation of C2C12 after 15d-PGJ_2_ treatment is rescued in C2C12 cells expressing the C-terminal mutants, specifically in cells expressing HRas-C184S (Fig. 2F and G). A previous study has demonstrated that 15d-PGJ_2_ covalently modifies Cys184 and not Cys181 in the C-terminal tail of HRas by means of the electrophilic center in the cyclopentenone ring(Luis Oliva et al., 2003). Therefore, the differentiation by C2C12 myoblasts expressing HRas-C184S but not by myoblasts expressing HRas WT after 15d-PGJ_2_ treatment suggests that 15d-PGJ_2_ inhibits differentiation of myoblasts by covalent modification of HRas at Cys184. We also show that 15d-PGJ_2_ preferentially activates the MAPK signaling pathway but not the PI3K signaling pathway, as seen by the phosphorylation of Erk and Akt proteins after 15d-PGJ_2_ treatment (Fig. 3A and B), even in the absence of growth factors. Both MAPK and PI3K signaling pathways have been reported to regulate muscle regeneration(Rommel et al., 1999). Therefore, preferential activation of the MAPK signaling pathway over the PI3K signaling pathway by activation of HRas could be a mechanism behind the inhibition of myoblast differentiation by 15d-PGJ_2_. We also observed a decrease in levels of HRas WT at the Golgi compared to the plasma membrane after 15d-PGJ_2_ treatment, unlike in the case of HRas C181S and C184S (Fig. 2A and B). This suggests that 15d-PGJ_2_ alters the intracellular distribution and cellular activity of HRas in a Cys184 and Cys181-dependent manner (Fig 3). We observed that the increase in the phosphorylation of Erk after 15d-PGJ_2_ treatment was diminished in C2C12 cells expressing HRas-C181S and not HRas-C184S even though 15d-PGJ_2_ has been shown to covalently modify HRas at Cys184(Luis Oliva et al., 2003). One possible reason behind this observation could be due to the reason that we overexpressed the HRas-C181S and HRas-C184S constructs in C2C12 cells which still had the basal expression of the endogenous HRas. Therefore, it is possible that 15d-PGJ_2_ could modify Cys184 of the endogenous HRas even in the presence of C184S HRas mutant and increase the overall phosphorylation of Erk. In case of HRas-C181S overexpression, it is expected that 15d-PGJ_2_ reacted with Cys184 of both the endogenous HRas and the overexpressed HRas-C181S. This could be the reason why Erk phosphorylation was not diminished in cells overexpressing the HRas-C184S mutant.

Senescent cells synthesize and secrete 15d-PGJ_2_ as part of the Senescence Associated Secretory Phenotype (SASP)(Martien et al., 2013; Ricciotti and Fitzgerald, 2011; Rossi et al., 1997; Wang and Dubois, 2006; Wiley et al., 2021). In this study, we show that the induction of senescence in C2C12 myoblasts by Doxo leads to the increased synthesis and secretion of 15d-PGJ_2_, as seen by the increase in the concentrations of 15d-PGJ_2_ both within senescent myoblasts and in their conditioned medium (Fig. 4A and B). We also show an increase in the expression of mRNA of prostaglandin biosynthetic enzymes (including PTGDS) in the hindlimb muscles of mice treated with Doxo (Fig. 4D). Doxo is a widely used anticancer drug(Johnson-Arbor and Dubey, 2022) and has been shown to induce senescence in cells(Di Leonardo et al., 1994; Hu and Zhang, 2019; Robles and Adami, 1998). Therefore, increased synthesis and secretion of prostaglandins is a possible mechanism underlying chemotherapy-induced loss of muscle mass and functioning(Yamaoka et al., 2015). Therefore, the targeted inhibition of only enzymes involved in the biosynthesis of specific prostaglandins which inhibit myoblast differentiation, e.g., PGD_2_ (via the enzyme PTGDS), will be less toxic than the pan-inhibition of all prostaglandin biosynthesis by NSAIDs. Targeting PTGDS using small molecules could be an important strategy to prevent the chemotherapy induces skeletal muscle loss in patients.

## Acknowledgements

We thank Prof. Satyajit Mayor (NCBS) and Prof. Apurva Sarin (InStem) for providing the wild-type HRas construct and vector backbones respectively. We thank Dr. Neetu Saini (InStem) for her help with setting up the cell culture facility. We thank Mr. Heera Lal for his help with the animal work. We thank Dr. Kamlesh Kumar Yadav and Ms. Sudeshna Saha for their help during the project. We thank the Central Imaging and Flow cytometry Facility (CIFF) (NCBS – InStem) for their support with confocal microscopy. We thank the Animal Care and Resource Center (NCBS – InStem) for their support with mouse experiments. We thank the Mass Spectrometry facility (NCBS – InStem) for their support with the mass spectrometry work.

## Author Contribution

Project Conceptualization: AR, SSP.

Animal work: AB.

Cell culture, assays, and treatments: SSP, SSS, AB, AV.

Biochemistry: SSP, SSS, AB.

Mass Spectrometry: AB, RGHM.

Microscopy, Image processing, and analysis: SSP, MAJ.

Writing – Original Draft: AR, SSP.

Writing – review, and editing: AR, SSP, AB, AV, SSS.

## Materials and Methods

### • Plasmids

Unmutated and cysteine mutants of HRas WT [HRas WT, HRas-C181S, and HRas-C184S] and HRas V12 [HRas V12, HRas V12-C181S, HRas V12-C184S] were cloned in the pEGFPC1 vector (Clontech) by restriction digestion-ligation method. Constructs of wild-type HRas were PCR amplified from a previously available HRas construct in the lab with construct-specific primers using Phusion High Fidelity DNA Polymerase (Thermo Scientific) followed by restriction digestion of HRas constructs and the pEGFPC1 vector using Xho1/EcoR1 enzymes (New England Biolabs Inc.). Ligation was set up using T4 DNA ligase (Takara Bio.). proper nucleotide additions were made to the forward primer to maintain the EGFP ORF, marking a 7 amino acid linker between the proteins. The construct sequences were confirmed by Sanger sequencing. GalT – TagRFP construct was a gift from Prof. Satyajit Mayor and was used to mark the Golgi.

### Cell Culture

#### Cell Maintenance

C2C12 mouse myoblasts (CRL – 1772) were obtained from ATCC and were maintained in DMEM complete medium @ 37° C, 5% CO_2_. For experiments, the cells were trypsinized with 0.25% trypsin – EDTA (Gibco): DPBS (Gibco) (1:1) @ 37° C, 3 mins. The cells were then resuspended in a triple volume of DMEM complete medium and were counted manually with a hemocytometer after diluting the cells with trypan blue (Gibco) (1:1) and were seeded in required numbers in cell culture dishes. Human Skeletal Muscle Myoblasts P2 (CC-2580) were obtained from Lonza and were maintained in DMEM Skeletal Muscle growth medium @ 37° C, 5% CO_2_. For experiments, the cells were trypsinized with 0.25% trypsin – EDTA (Gibco): DPBS (Gibco) (1:1) @ 37° C, 2 mins. The cells were then resuspended in a double volume of DMEM complete medium and were counted manually with a hemocytometer after diluting the cells with trypan blue (Gibco) (1:1) and were seeded in required numbers in cell culture dishes. All cultures tested negative for mycoplasma checked by Mycoalert Mycoplasma Detection Kit (Lonza).

#### Conditioned medium collection

3×10^5^ senescent cells/well or 1.5×10^5^ proliferative cells/well were seeded into 6-well plates. After overnight incubation, the media was changed to 2ml of 0.2% FBS containing DMEM. Conditioned medium was collected after 24 hr and the cells were trypsinized, counted, and pelleted down. The conditioned media was centrifuged @ 2000 rpm for 10 min to clear out debris, the supernatant was collected in a fresh tube. The supernatant and the cell pellet were flash-frozen in liq N_2_ and were stored at −80° C until analysis.

#### Treatments

##### 15d-PGJ_2_

15d-PGJ_2_ (Cayman Chemical Company) dissolved in methyl acetate was purged with N_2_ stream @ R.T. till drying and was then redissolved in DMSO to make a 10 mM stock solution. 15d-PGJ_2_ (10 mM) stock was then appropriately diluted in an appropriate volume of DMEM media for experiments. DMSO was used as vehicle control. A media change of the same composition was given every 24 hours.

##### 9,10-dihydro-15d-PGJ_2_

9,10-dihydro-15d-PGJ_2_ (Cayman Chemical Company) dissolved in methyl acetate was purged with N_2_ stream @ R.T. till drying and was then redissolved in DMSO to make a 10 mM stock solution. 9,10-dihydro-15d-PGJ_2_ (10 mM) stock was then appropriately diluted in an appropriate volume of DMEM media for experiments. DMSO was used as a control.

##### Doxorubicin

C2C12 cells with 70-80% confluency in a 10 cm plate were treated with Doxorubicin (Doxo) for 3 days. After 72hr, Doxo was removed from the medium and the cells were kept for 10 more days with media change every 3 days. The cells were harvested on day 13 cells after Doxo treatment for experiments.

#### Transfections

C2C12 cells were seeded in 35 mm dishes at an intermediate density to achieve confluency of ∼60-70%. For western blot and differentiation experiments, the cells were transfected with 1.5 μg of EGFP-tagged HRas WT/ HRas-C181S/ HRas-C184S/ HRas V12/ HRas V12-C181S/ HRas V12-C184S using the jetPRIME transfection reagent (Polyplus) using the manufacturer’s protocol. Transfection efficiency was confirmed by checking for GFP fluorescence in cells under a Ti2 epifluorescence microscope (Nikon) using appropriate filters. For measuring the intracellular distribution of HRas, the cells were reverse transfected with 1μg each of EGFP-tagged HRas WT/ HRas-C181S/ HRas-C184S/ HRas V12/ HRas V12-C181S/ HRas V12-C184S and GalT - TagRFP, a Golgi apparatus marker protein tagged with red fluorescent TagRFP protein using jetPRIME, where the DNA - jetPRIME mixture was incubated with the cell suspension while seeding the cells after trypsinization, and the transfection efficiency was confirmed by checking GFP and RFP fluorescence after 24 hours of transfection.

#### Myoblast differentiation

##### Untransfected cells

C2C12 cells were seeded in 35 mm dishes (Corning) at high density to achieve a confluency of ∼90 – 95% the next day. The cells were then treated with either 15d-PGJ_2_ or DMSO in the C2C12 differentiation medium. The cells were given a media change of the same composition every 24 hours. The cells were harvested after 5 days of 15d-PGJ_2_ treatment for either RNA or protein isolation. For the Immunofluorescence experiment, the experiment was done in 35 mm dishes (Corning) on glass coverslips (Blue star) coated with 0.2% Gelatin (Porcine, Sigma Aldrich), and the cells were fixed with the fixative solution at the end of the experiment and were immunostained. Human Skeletal Muscle Myoblast cells were seeded in 35 mm dishes in high density to achieve a confluency of ∼90 – 95% the next day. The cells were then treated with DMSO or 15d – PGJ_2_ in the Skeletal Muscle Differentiation medium. A media change of the same composition was given every 24 hours. The cells were harvested after 5 days of treatment in RIPA - PP.

##### Transfected cells

C2C12 cells were seeded in 35 mm dishes (Corning) at an intermediate density to achieve a confluency of ∼60 – 70% confluency. The cells were transfected with EGFP-tagged HRas WT/ HRas-C181S/ HRas-C184S/HRas V12/HRas V12-C181S/ HRas V12-C184S using jetPRIME transfection reagent (Polyplus). After confirming a transfection efficiency of ∼80% the next day, the cells were treated with DMSO vehicle or 15d – PGJ_2_ (4 µM) in the C2C12 differentiation medium. The cells were given a media change of the same composition every 24 hours. The cells were harvested after 5 days of 15d – PGJ_2_ treatment for either RNA or protein isolation.

#### X-Gal staining

Proliferative and day-13 dox-treated cells were seeded at a density of 1×10^5^ cells /well into 35mm dishes. After overnight incubation, SA-β-gal activity was measured by using Senescence β-Galactosidase Staining Kit (Cell Signalling #9860) per manufacture’s protocol. In brief, the cells were washed with PBS twice and were fixed for 10-15 min at RT. Following three washes of PBS, cells were incubated overnight in staining solution at 37℃ in a CO_2_–free chamber. Development of blue colour was examined with Ti2 widefield inverted microscope (Nikon).

### Gene expression analysis

#### Western blotting

For measuring Erk/Akt phosphorylation in C2C12 cells (untransfected/ transfected with EGFP-tagged HRas WT/ HRas-C181S/ HRas-C184S/ HRas V12/ HRas V12-C181S/ HRas V12-C184S) were seeded in 35 mm dishes. 1x 35 mm dish was harvested in RIPA – PP the next day, while the rest were incubated in DMEM starvation medium @37° C. The cells were treated with 15d - PGJ_2_ after 24 hrs of starvation @37° C. The cells were harvested 1 hr after treatment in RIPA - PP. Protein quantification was done using BCA assay (G Biosciences) using the manufacturer’s protocol. An equal mass of proteins was loaded onto a 12% SDS Polyacrylamide gel in Laemmlli buffer. The proteins were transferred onto a PVDF membrane and were probed with phospho-Erk/Erk antibodies for measuring Erk phosphorylation and with phospho-Akt/Akt antibodies for measuring Akt phosphorylation. For measuring the expression of Myosin heavy chain, C2C12 cells expressing EGFP-tagged HRas WT/ HRas-C181S/ HRas-C184S/ HRas V12/ HRas V12-C181S/ HRas V12-C184S or Human Skeletal Muscle Myoblasts were seeded in 35 mm dishes and were harvested in RIPA - PP after 5 days of differentiation. Protein quantification was done using BCA assay (G Biosciences) using the manufacturer’s protocol. An equal mass of proteins was loaded onto an 8% SDS Polyacrylamide gel in Laemmlli buffer. The proteins were transferred onto a PVDF membrane and were probed with Myosin Heavy Chain Antibody.

#### qPCR

C2C12 cells, untransfected or expressing EGFP-tagged HRas WT/ HRas-C181S/ HRas-C184S and treated with DMSO/15d - PGJ_2_, or expressing EGFP-tagged HRas V12/ HRas V12-C181S/ HRas V12-C184S were lysed in TRIZol at the end of the experiment (Invitrogen). RNA was isolated from the lysate by the chloroform-isopropanol method using the manufacturer’s protocol. The RNA was quantified and 1.5 μg of RNA was used to prepare cDNA using PrimeScript 1st strand cDNA Synthesis Kit (Takara Bio) and random hexamer primer. Gene expression for differentiation markers was measured by qPCR using PowerUp™ SYBR™ Green Master Mix (Applied Biosystems) and previously reported qPCR primers (p.m. 43). Relative gene expression was quantified using the ΔΔC_T_ method^44^ with 18s rRNA as an internal loading control and DMSO vehicle as an experimental control.

#### Immunofluorescence

C2C12 cells were seeded in 35 mm dishes (Corning) on glass coverslips (Blue Star) coated with 0.2% Gelatin (Porcine, Sigma Aldrich) and were fixed with the fixative solution at the end of the experiment. The cells were then permeabilized and blocked with the blocking solution and were then incubated with MyHC antibody in the blocking solution overnight. The cells were then washed with 1x PBS, incubated with fluorophore tagged secondary antibody, and were mounted in Prolong gold antifade medium with DAPI (Invitrogen). The cells were then imaged under FV3000 inverted confocal laser scanning microscope (Olympus – Evident) using appropriate lasers and detectors.

### HRas distribution between the Golgi and the plasma membrane

C2C12 cells expressing EGFP-tagged HRas WT/ HRas-C181S/ HRas-C184S + GalT-TagRFP were starved overnight in DMEM starvation medium and treated with DMSO vehicle or 15d – PGJ_2_ (10 µM) in DMEM complete medium for 24 hrs, with a medium change @ 12 hrs post-treatment. The cells were then fixed with the fixative solution @ R. T., 5 mins, washed with PBS, and stained for plasma membrane with Alexa Fluor 633 conjugated Wheat Germ Agglutinin (WGA - 633) (Invitrogen) (10 µg/ml) @R. T. 15 mins. The cells were washed with PBS and were then mounted on glass slides in ProLong Gold Antifade Mounting medium (Invitrogen). C2C12 cells expressing EGFP-tagged HRas V12/ HRas V12-C181S/ HRas V12-C184S were also fixed with the fixative solution @R. T., 5 mins, washed with PBS, stained with WGA – 633, and mounted on slides in Prolong gold antifade medium (Thermo Scientific). The cells were imaged with FV 3000 inverted confocal laser scanning microscope (Olympus - Evident) using appropriate lasers and detectors. Preliminary image processing was done using ImageJ (NIH), while batch analysis of HRas at the plasma membrane and the Golgi complex was done using a custom MATLAB script, where EGFP-HRas image was overlayed onto the GalT-TagRFP and WGA-633 image to obtain HRas localization at the Golgi complex and the Plasma Membrane respectively. A ratio of mean HRas intensity at the Golgi complex to that of at the Plasma membrane (R_mean_) was calculated and was used to compare HRas distribution between treatments.

### Animal care

#### Maintenance

Mice were maintained at BLiSC Animal Care and Resource Centre (ACRC). All the procedures performed were approved by the Internal Animal Users Committee (IAUC) and the Institutional Animal Ethics Committee (IAEC).

#### Treatment and tissue collection

12–15-week-old C57BL/6J (JAX#000664) mice were injected intraperitoneally (I.P.) with 5 mg/kg Doxorubicin (Doxo) four times, once every three days. Intraperitoneal injection of Saline was used as a control. The mice were sacrificed on Day 11 after the first injection. Hindlimb muscles from 4 animals (control and treated with Doxo each) were used for qPCR analysis and Hindlimb muscles from 3 animals (control and treated with Doxo each) were used for immunohistochemical analysis.

### Lipid extraction and detection of 15d-PGJ_2_ by mass spectrometry

For lipid extraction, cell pellets were resuspened in 3ml of a methanol solvent [water: methanol: 2:1, 1% formic acid (FA)] whereas only 1 ml of methanol with 3% FA was added to the 2ml of CM, making a uniform sample volume of 3 ml. Subsequently, 1ml of ethyl acetate was added to each sample and mixed vigorously. Phase separation was done by centrifuging the mixture (12000xg, 4℃ for 10 mins), and the organic phase containing the lipid was collected. This process was repeated thrice in total and all the organic phases were combined and dried under a nitrogen stream at RT. The residues were resuspended in 100 µl of 50% acetonitrile in water with 0.1% FA and were subjected to mass spec analysis using the Waters® Acquity UPLC class I system The detection of 15d-PGJ_2_ was performed using an electrospray ionization source (ESI) operating in the negative ion mode and a quadrupole trap mass spectrometer (AB SCIEX QTRAP 6500) connected to a Waters® Acquity UPLC class I system (Waters, Germany) outfitted with a binary solvent delivery system with an online degasser and a column manager with a column oven coupled to a UPLC autosampler. 5 µl samples were injected into the union for analysis. Solvent A consisted of 0.1% ammonium acetate in water and solvent B was 0.1% ammonium acetate in a mixture of acetonitrile/water (95:5). For each run, the LC gradient was: 0 min, 20% B; 0.5 min, 20% B; 1.5 min, 90% B; 2.5 min, 20% B; 3min, 20% B. Analyte detection was performed using multiple reaction monitoring (MRM), 315.100 ➔ 271.100 and 315.100 ➔ 203.100. Source parameters were set as follows: capillary voltage 3.8 kV, desolvation gas flow 25 L/h, source temperature 350 °C, ion source gas 1 flow 40 L/h and ion source gas 2 flow 40 L/h. Acquisition and quantification were completed with Analyst 1.6.3 and Multiquant 3.0.3, respectively (method adopted from (Morgenstern et al., 2018)).

For the standards, 2ml media of different known concentrations (50nM, 100nM, 250nM, and 500nM) of 15d-PGJ_2_ were prepared and were subjected to the same extraction procedure as that of CM. A standard curve was plotted with the known concentration and the mass spec peak area, and the concentration of the lipid in samples was calculated.

**Table 1:**
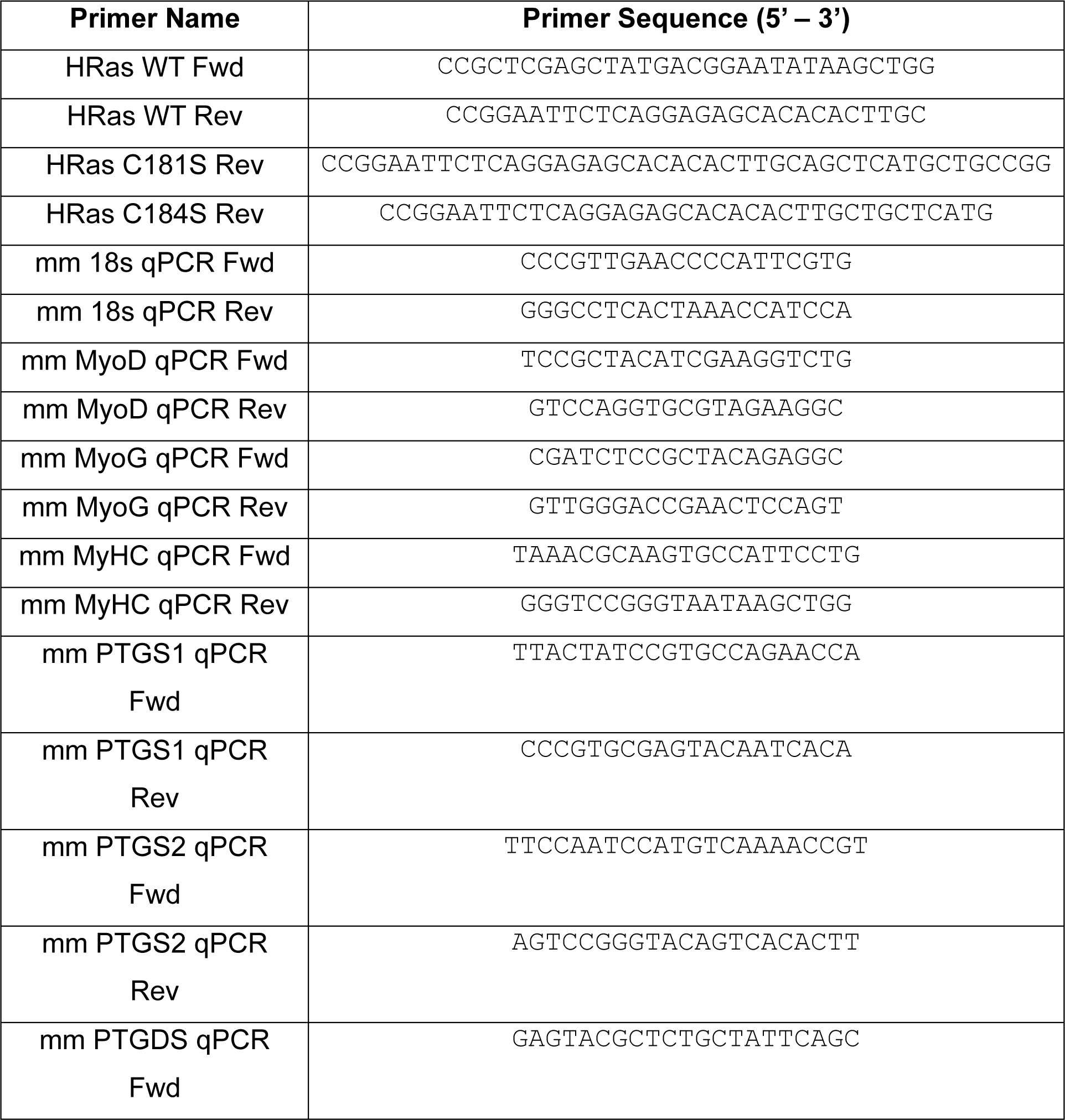
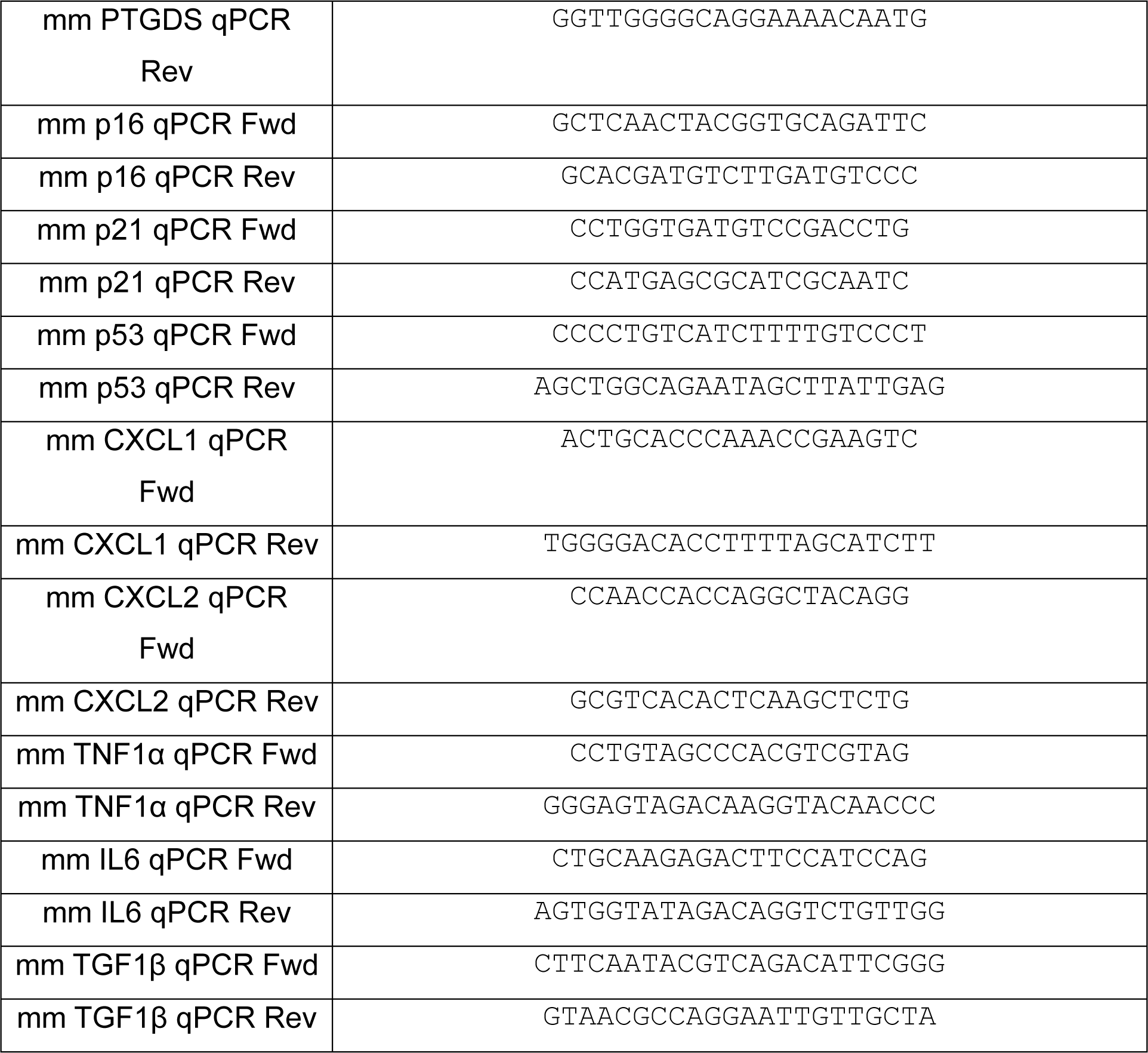
List of Primers:

**Table 2:** List of Reagents

| Item Description | Manufacturer | Catalog No. |
| --- | --- | --- |
| pEGFP-C1 | Clontech |  |
| Phusion high fidelity DNA polymerase | Thermo Scientific | F 530 |
| Xho1 restriction enzyme | New England Biolabs Inc. | R0146 |
| EcoR1 restriction enzyme | New England Biolabs Inc. | R0101 |
| T4 DNA Ligase | Takara Bio | 2011 |
| DMEM High Glucose | Gibco | 11995065 |
| FBS, Certified (Origin: United States) | Gibco | 16000044 |
| Horse Serum, Heat Inactivated (Origin: New Zealand) | Gibco | 26050070 |
| Penicillin – Streptomycin – Glutamine (100x) | Gibco | 10378016 |
| Penicillin – Streptomycin (100x) | Gibco | 15140163 |
| DPBS, no calcium, no magnesium | Gibco | 14190144 |
| 0.25% Trypsin - EDTA | Gibco | 25200056 |
| jetPRIME transfection reagent | Polyplus | 101000046 |
| Doxorubicin | Sigma – Aldrich | D1515 |
| 15-deoxy-$\Delta^{12,14}$-Prostaglandin J<sub>2</sub> | Cayman Chemical Company | 18570 |
| 9,10-dihydro-15-deoxy-$\Delta^{12,14}$-Prostaglandin J<sub>2</sub> | Cayman Chemical Company | 18590 |
| cOmplete Protease inhibitor cocktail tablets | Roche | 11697498001 |
| Wheat germ agglutinin, Alexa fluor 633 conjugate | Invitrogen | 2307289 |
| ProLong Gold Antifade Mounting medium | Invitrogen | P36930 |
| Paraformaldehyde | Sigma – Aldrich | 158127 |
| TRIzol reagent | Invitrogen | 15596018 |
| PrimeScript™ 1st strand cDNA Synthesis Kit | Takara Bio | 6110A |
| PowerUp™ SYBR™ Green Master Mix | Applied Biosystems | A25742 |
| WesternBright™ ECL-spray Western blotting detection system | advansta | K-12049-D50 |
| Fetuin (Bovine) | Sigma – Aldrich | F2379 |
| hEGF | Sigma – Aldrich | E9644 |
| N2 Supplement | Thermo Scientific | 17502048 |
| Dexamethasone | Sigma – Aldrich | D4902 |
| DMEM Low Glucose | Thermo Scientific | 10567014 |
| Bovine Serum Albumin Fraction-V, Cell culture tested | HIMEDIA | 9048468 |
| Pierce™ Streptavidin Magnetic Beads | Thermo Scientific | 88816 |

**Table 3:** List of Antibodies

| Antibody Name | Clonality | Species of Origin | Manufacturer | Catalog No. |
| --- | --- | --- | --- | --- |
| Phospho-Erk (Thr202/<br>Tyr204) | Polyclonal | Rabbit | Cell Signaling Technology | 9101S |
| Erk | Polyclonal | Rabbit | Cell Signaling Technology | 9102S |
| GAPDH | Monoclonal | Mouse | Puregene | PG23002 |
| β-actin | Polyclonal | Rabbit | Cell Signaling Technology | 4967S |
| Phospho-Akt (Ser473) | Monoclonal | Mouse | Cell Signaling Technology | 4051S |
| Akt | Monoclonal | Rabbit | Cell Signaling Technology | 4691S |
| GFP | Monoclonal | Rabbit | Cell Signaling Technology | 2956S |
| Myosin Heavy Chain | Monoclonal | Mouse | Invitrogen | 14-6503-82 |
| p21 | Monoclonal | Mouse | Santacruz Biotechnology | sc-6246 |
| γH2A.X | Polyclonal | Rabbit | Novus Biologicals | NB100384 |
| Tubulin | Monoclonal | Mouse | Cell Signaling Technology | 3873S |
| HRP - Anti-Mouse |  | Horse | Cell Signaling Technology | 7076S |
| HRP - Anti-Rabbit |  | Goat | Cell Signaling Technology | 7074P2 |
| Alexa Fluor 568 – Anti-Mouse | Polyclonal | Goat | Invitrogen | A11031 |

## Reagent Composition

- DMEM complete medium: DMEM Hi Glucose medium (Gibco) supplemented with 1% Penicillin – Streptomycin – Glutamine (Gibco) and heat-inactivated 10% Fetal Bovine Serum (US origin) (Gibco).
- Basal Conditioned medium: DMEM Hi Glucose medium (Gibco) supplemented with 1% Penicillin – Streptomycin – Glutamine (Gibco) and heat-inactivated 2% Fetal Bovine Serum (US origin) (Gibco).
- C2C12 differentiation medium: DMEM Hi Glucose medium (Gibco) supplemented with 2% Horse Serum (Gibco) and 1% Penicillin – Streptomycin – Glutamine (Gibco).
- DMEM Starvation medium: DMEM Hi Glucose medium (Gibco) supplemented with 0.2%heat-inactivated fetal bovine serum (US origin) (Gibco) and 1% Penicillin – Streptomycin – Glutamine (Gibco).
- RIPA – PP buffer: RIPA buffer (Invitrogen) supplemented with protease inhibitor cocktail (Roche) and 5 mM Sodium Fluoride and 5 mM Sodium Orthovanadate.
- TBS – T buffer: 50 mM Tris–Cl (pH = 7.5), 150 mM NaCl and 0.1% Tween – 20 in water.
- PBS: 2.67 mM KCl, 1.47 mM KH_2_PO_4_, 137.93 mM NaCl, 8.06 mM Na_2_HPO_4_ in water.
- Fixative Solution: 4% (w/v) Paraformaldehyde (Sigma – Aldrich) in PBS.
- Blocking Solution: 2% Heat Inactivated FBS, 0.2% BSA, 0.2% Triton - X, 0.05% NaN_3_ in PBS.
- Skeletal Muscle Growth Medium: DMEM Low Glucose Medium (Gibco), supplemented with 1% Penicillin – Streptomycin – Glutamine (Gibco), heat-inactivated 10% Fetal Bovine Serum (US origin) (Gibco), Bovine Fetuin (50 µg/ml) (Sigma – Aldrich), Dexamethasone (0.4 µg/ml), and hEGF (10 ng/ml).
- Skeletal Muscle Differentiation Medium: DMEM low glucose medium (Gibco) supplemented with 2% Horse Serum, 1% Penicillin – Streptomycin (Gibco), and 1% N2 Supplement.

