## Supplementary figures for "C-terminal cysteines of HRas control Erk signaling and 15-deoxy-Δ^12,14^-prostaglandin J^2^ (15d-PGJ^2^) mediated inhibition of myoblast differentiation"

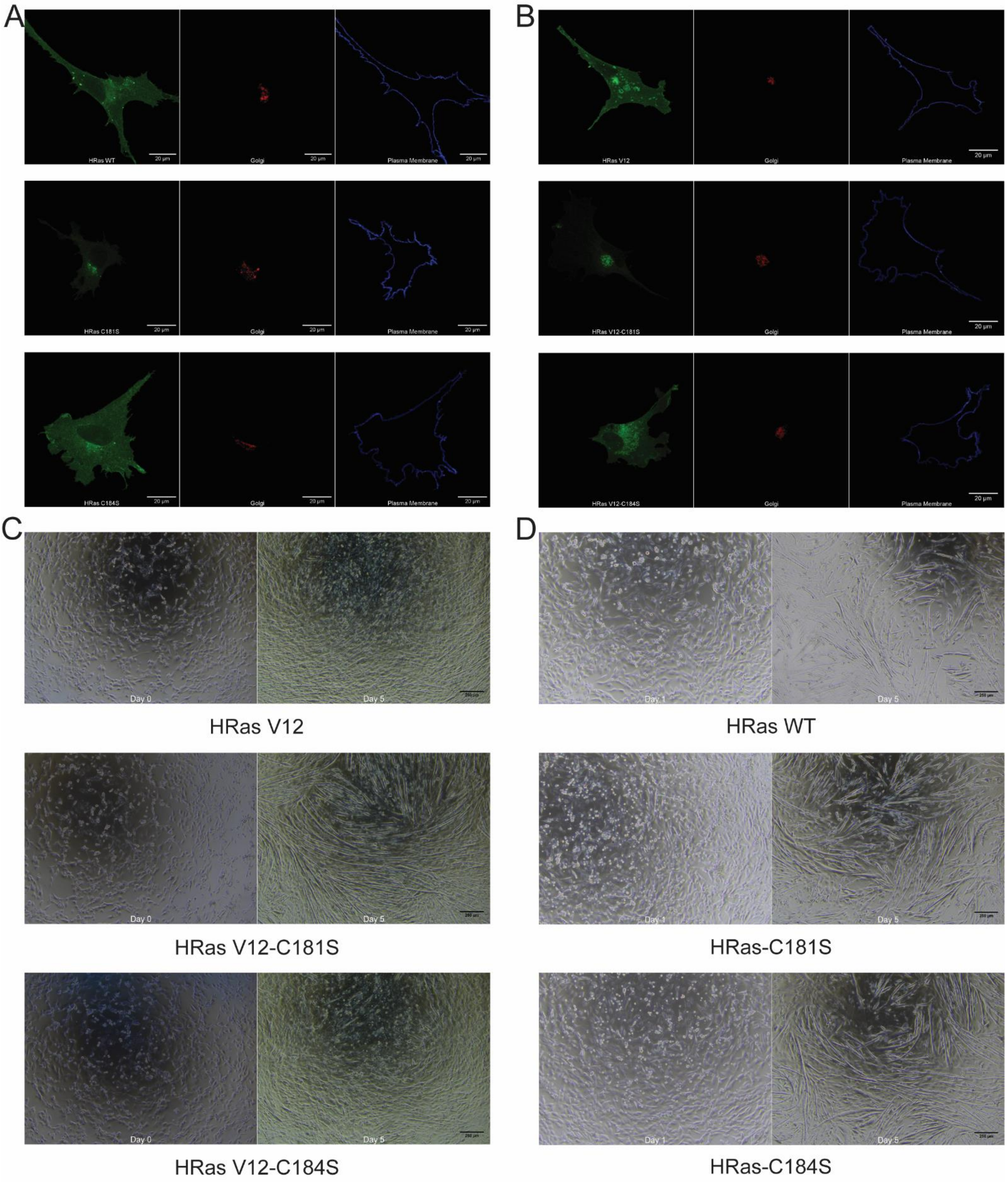


**Supplementary Figure S1: The effect of C-terminal cysteines of HRas on its intracellular distribution and differentiation of C2C12 myoblasts.**

A. A representative fluorescent confocal micrograph of C2C12 cell showing localization of HRas WT or HRas-C181S or HRas-C184S (EGFP-tagged) between the plasma membrane (stained with Alexa Fluor 633 conjugated Wheat Germ Agglutinin) and the Golgi complex (tagged with TagRFP conjugated Golgi resident GalT protein).
B. A representative fluorescent confocal micrograph of C2C12 cell showing localization of HRas V12 or HRas V12-C181S or HRas V12-C184S (EGFP-tagged) between the plasma membrane (stained with Alexa Fluor 633 conjugated Wheat Germ Agglutinin) and the Golgi complex (tagged with TagRFP conjugated Golgi resident GalT protein).
C. A representative bright field image of differentiating C2C12 cells expressing EGFP-tagged HRas V12 or HRas V12-C181S or HRas V12-C184S.
D. A representative bright field image of differentiating C2C12 cells expressing EGFP-tagged HRas WT or HRas-C181S or HRas-C184S.

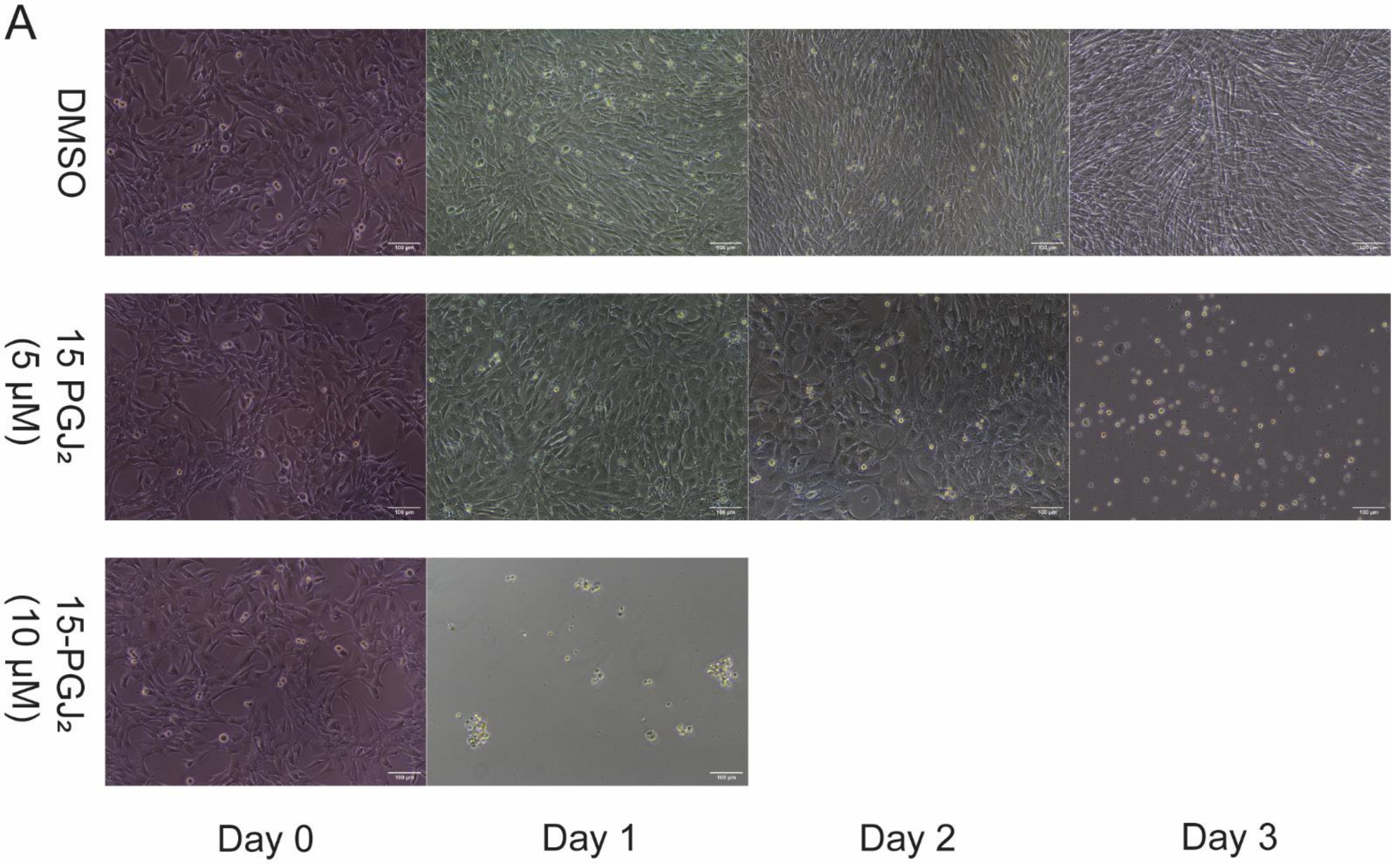


**Supplementary Figure S2: Reduced viability of C2C12 myoblasts after treatment with 15d-PGJ$_2$ (5 and 10 µM) in differentiating medium.**

A. Representative images taken every 24 hrs of C2C12 treated with 15d-PGJ$_2$ (5 and 10 µM) or DMSO in the C2C12 differentiation medium.

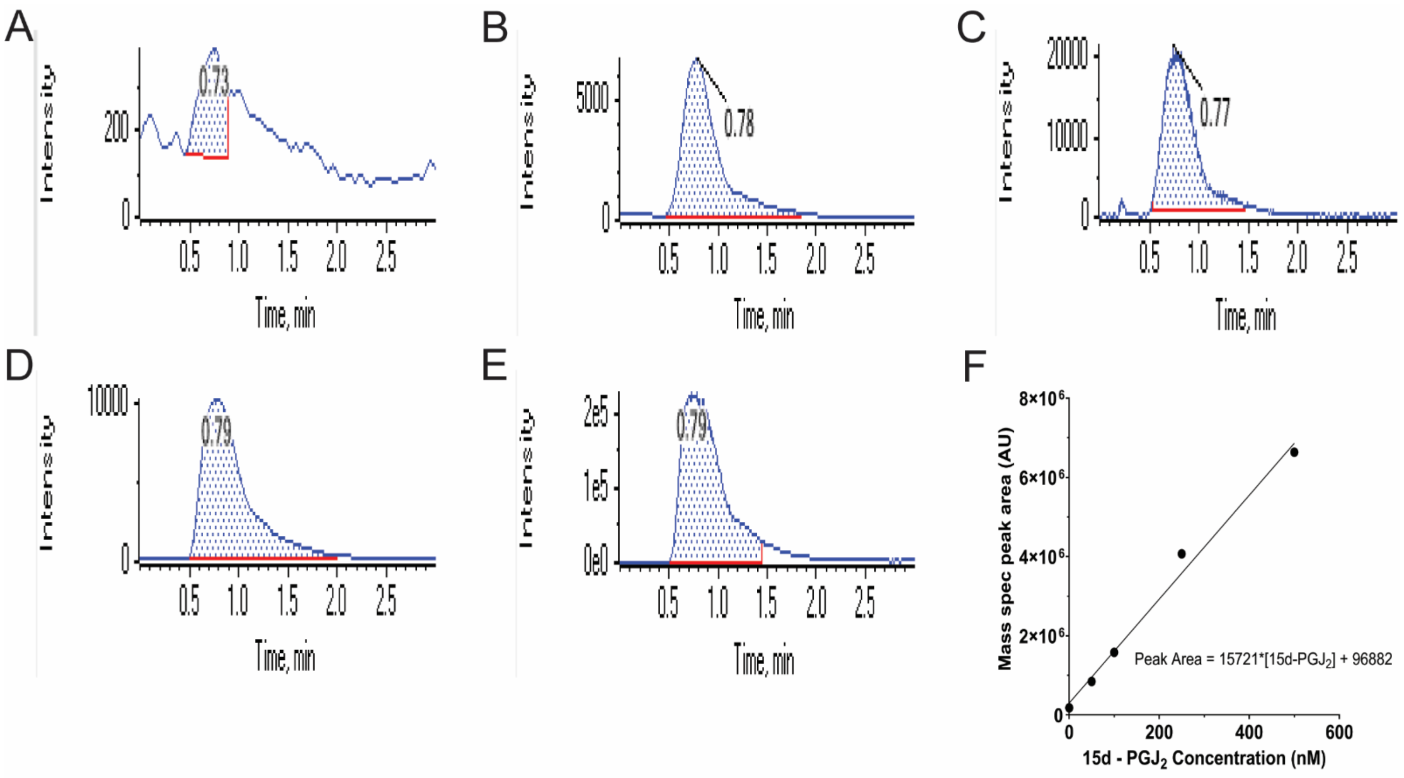


**Supplementary Figure S3: Quantification of intracellular 15d-PGJ$_2$ concentration and concentration of 15d-PGJ$_2$ from conditioned medium of C2C12 myoblasts treated with Doxo (150 nM) or DMSO.**

A. Representative peaks from Blank sample.
B. Representative peaks from C2C12 myoblasts treated with DMSO.
C. Representative peaks from C2C12 myoblasts treated with Doxo (150 nM).
D. Representative peaks from Conditioned medium of C2C12 myoblasts treated with DMSO.
E. Representative peaks from Conditioned medium of C2C12 myoblasts treated with Doxo (150 nM).
F. Standard curve of peak area vs known conc. of 15d-PGJ$_2$ detected using AB SCIEX QTRAP 6500 mass spectrometer.

| Sample | 15d-$PGJ_2$ (pmol) | Cell Number | 15d-$PGJ_2$/Cell (fg) |
|---|---|---|---|
| **Intracellular Quiescent C2C12 cells_1** | 27.087 | 462500 | 18.531 |
| **Intracellular Quiescent C2C12 cells_1** | 18.080 | 650000 | 8.801 |
| **Intracellular Senescent C2C12 cells_1** | 809.633 | 287500 | 891.018 |
| **Intracellular Senescent C2C12 cells_1** | 940.032 | 300000 | 991.420 |
| **Conditioned medium Quiescent C2C12 cells_1** | 8.271 | 462500 | 5.659 |
| **Conditioned medium Quiescent C2C12 cells_2** | 6.554 | 650000 | 3.190 |
| **Conditioned medium Senescent C2C12 cells_1** | 45.814 | 287500 | 50.419 |
| **Conditioned medium Senescent C2C12 cells_2** | 38.893 | 300000 | 41.019 |

**Supplementary Table 1: 15d-$PGJ_2$ conc. (pmol and fg/cell) detected in samples by mass spectrometry.**

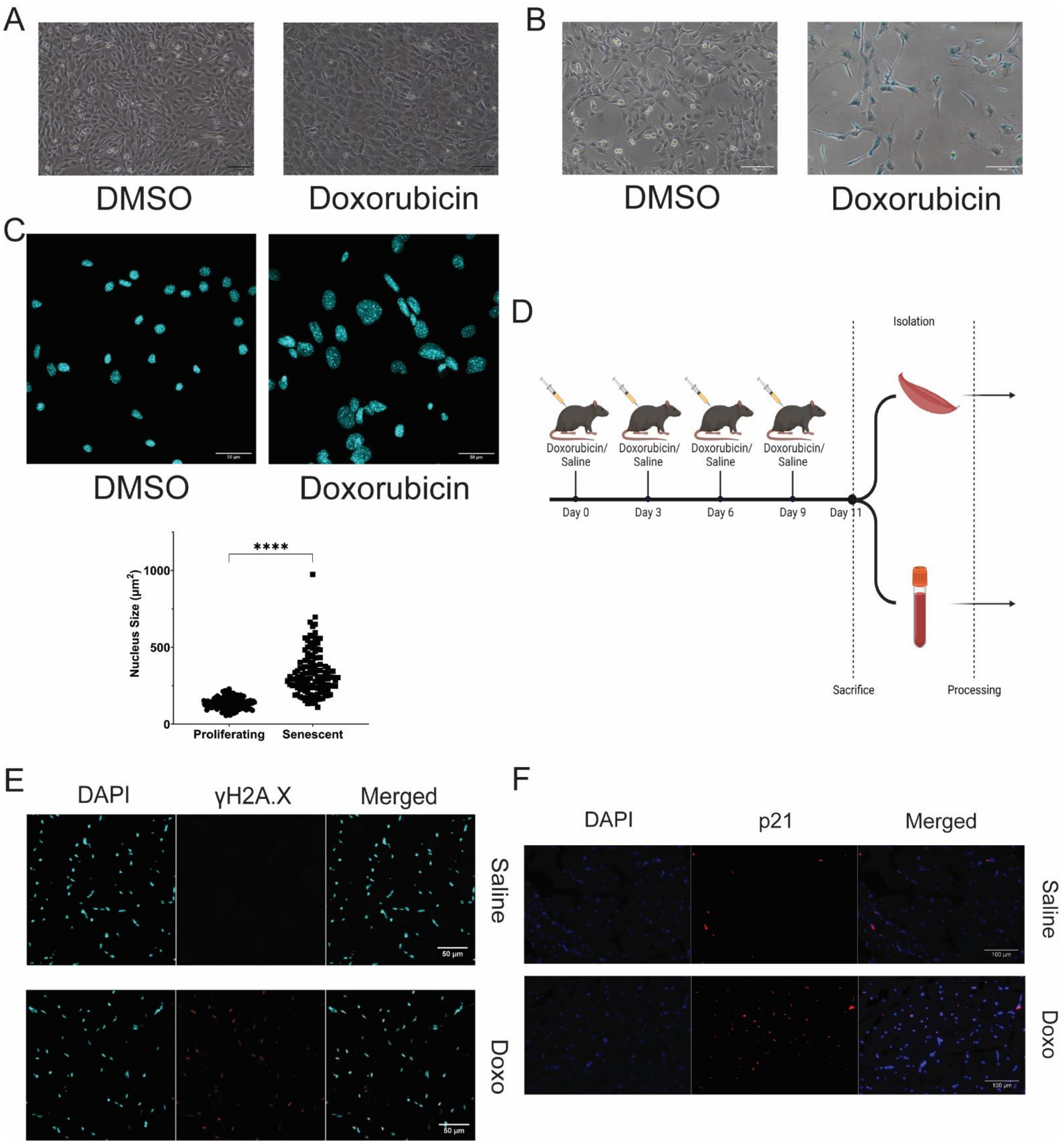


**Supplementary Figure S4: Induction of senescence after Doxorubicin (Doxo) treatment in C2C12 myoblasts and B6J mice.**

A. A representative bright field image of C2C12 cells after 5 days of treatment with Doxo (150 nM) or DMSO.
B. X-gal staining of C2C12 cells after 13 days of treatment with Doxo (150 nM) or DMSO showing SA-β gal activity.
C. A representative confocal micrograph and a scatter plot of nuclear size in C2C12 cells after 13 days of treatment with Doxo (150 nM) or DMSO.

D. Schematic representation of treatment of B6J mice with Doxo (5mg/kg) or Saline for induction of senescence.
E. A representative confocal micrograph of γH2A.X expression in the gastrocnemius muscle of B6J mice treated with Doxo or Saline.
F. A representative widefield epifluorescence image of p21 nuclear localization in the gastrocnemius muscle of B6J mice treated with Doxo or Saline.
